# The tumor suppressor APC is an attenuator of spindle-pulling forces during *C. elegans* asymmetric cell division

**DOI:** 10.1101/157404

**Authors:** Kenji Sugioka, Lars-Eric Fielmich, Kota Mizumoto, Bruce Bowerman, Sander van den Heuvel, Akatsuki Kimura, Hitoshi Sawa

**Affiliations:** Multicellular Organization Laboratory, National Institute of Genetics, 1111 Yata, Mishima, 411-8540 Japan; RIKEN Center for Developmental Biology, 2-2-3 Minatojima-minamimachi, Chuoku, Kobe 650-0047 Japan; Institute of Molecular Biology, University of Oregon, Eugene, OR 97403 USA; Developmental Biology, Biology Department, Science 4 Life, Utrecht University, Padualaan 8, 3584 CH, Utrecht, Netherlands; Cell Architecture Laboratory, National Institute of Genetics, 1111 Yata, Mishima, 411-8540 Japan; Department of Genetics, School of Life Science, Sokendai, 1111 Yata, Mishima, 411-8540 Japan

**Author notes:** Hitoshi Sawa, Multicellular Organization Laboratory, National Institute of Genetics, 1111 Yata, Mishima, 411-8540 Japan. Akatsuki Kimura, Cell Architecture Laboratory, National Institute of Genetics, 1111 Yata, Mishima, 411-8540 Japan. Sander van den Heuvel, Developmental Biology, Biology Department, Utrecht University, Padualaan 8, 3584 CH, Utrecht, Netherlands. Present address: Kota Mizumoto, Department of Zoology, the University of British Columbia, Vancouver, Canada, V6T 1Z3.

## Abstract

The adenomatous polyposis coli (APC) tumor suppressor has dual functions in Wnt/ß-catenin signaling and accurate chromosome segregation, and is frequently mutated in colorectal cancers. Although APC contributes to proper cell division, the underlying mechanisms remain poorly understood. Here we show that *C. elegans* APR-1/APC is an attenuator of the pulling forces acting on the mitotic spindle. During asymmetric cell division of the *C. elegans* zygote, a LIN-5/NuMA protein complex localizes dynein to the cell cortex to generate pulling forces on astral microtubules that position the mitotic spindle. We found that APR-1 localizes to the anterior cell cortex in a Par-aPKC polarity-dependent manner and suppresses anterior centrosome movements. Our combined cell biological and mathematical analyses support the conclusion that cortical APR-1 reduces force generation by stabilizing microtubule plus ends at the cell cortex. Furthermore, APR-1 functions in coordination with LIN-5 phosphorylation to attenuate spindle pulling forces. Our results document a physical basis for spindle-pulling force attenuation, which may be generally used in asymmetric cell division, and when disrupted potentially contributes to division defects in cancer.

**Significance Statement:** APC (adenomatous polyposis coli) is a Wnt signaling component as well as a microtubule-associated protein, and its mutations are frequently associated with colorectal cancers in humans. Although APC stabilizes microtubules (MTs), its mechanical role during cell division is largely unknown. Here we show that APC is an attenuator of forces acting on the mitotic spindle during asymmetric cell division of the *C. elegans* zygote. We performed live-imaging, laser-microsurgery, and numerical simulation to show how APC suppresses spindle pulling force generation by stabilizing microtubule plus-ends and reducing microtubule catastrophe frequency at the cell cortex. Our study is the first to document a mechanical role for the APC protein, and provides a physical basis for spindle-pulling force attenuation.

## Introduction

The mitotic spindle segregates chromosomes and determines the plane of cell cleavage during animal cell division. Forces that act on the mitotic spindle regulate its position to produce daughter cells of the proper size, fate and arrangement, thereby playing a significant role in asymmetric cell division, tissue integrity and organogenesis. In various organisms, cells regulate spindle positioning through cortical force generators that pull on astral microtubules (1-5). An evolutionarily conserved force generator complex, consisting of LIN-5/NuMA, GPR-1,2/LGN and Gα, interacts with dynein and dynamic astral microtubules to position the mitotic spindle during the asymmetric divisions of the *C. elegans* early embryo (4), *Drosophila* and mammalian neuroblasts (1, 2), and skin stem cells (3). Although Par-aPKC polarity and cell cycle regulators are known to control spindle positioning (4, 6), how the forces are regulated spatiotemporally to position the spindle in various cell types during development remains poorly understood.

The tumor suppressor adenomatous polyposis coli (APC) is a widely conserved multifunctional protein with two major roles. First, APC functions as part of a degradation complex to down-regulate β-catenin-TCF dependent transcription, thereby controlling cell fate and proliferation in various cell types (7). Second, APC functions as a microtubule-associated protein to stabilize MTs. It has been suggested that this function of APC regulates cell migration (8, 9), spindle orientation (10, 11), and chromosome segregation (12, 13). In mammals, loss of the former function is closely associated with colon cancer (14, 15). Loss of the latter function causes spindle positioning defects (16, 17) and chromosome instability (CIN) (18-20), a hallmark of metastatic tumors (21), suggesting that the cytoskeletal roles of APC during mitosis are also relevant for oncogenesis. How APC regulates the mitotic spindle remains poorly understood and is complicated by its multiple functions, binding-partners and cellular locations (12, 22).

Yeast and fly studies have suggested that APC at the cell cortex contributes to mitotic spindle positioning. Kar9, a yeast protein with limited homology to APC, localizes asymmetrically to the cell cortex of budding daughter cells through type V myosin-dependent transport of growing microtubule ends (23-25). Cortical Kar9 captures microtubules (MTs) by binding yeast EB1, and promotes alignment of the spindle along the mother-bud axis (24-27). *Drosophila* APC2 predominantly localizes to the cell cortex in syncytial embryos. APC2 mutants show a CIN phenotype, presumably because APC2 is required for proper centrosome separation (28). The forces that mediate centrosome separation have been proposed to depend on APC2 connecting astral MTs to cortical actin (28). However, the mechanism by which cortical APC regulates spindle-pulling forces has not been directly addressed in any organism.

We report here that loss of cortical APR-1/APC disrupts asymmetries in spindle movements during mitotic division of the *C. elegans* zygote. In wild-type embryos, the net pulling forces acting on the mitotic spindle become higher in the posterior compared to the anterior, causing the spindle to move posteriorly during metaphase and anaphase (spindle displacement) (29, 30). In anaphase, the posterior spindle pole swings along the transverse axis (spindle oscillation), while the anterior pole remains relatively stable. We found APR-1 to be enriched at the anterior cortex in a PAR-polarity dependent manner. Depletion of APR-1 resulted in anterior pole oscillations that resemble those of the posterior pole. Moreover, laser-mediated spindle severing showed that the spindle-pulling forces acting on the anterior spindle pole are increased in *apr-1(RNAi)* embryos. Using live imaging and numerical simulation, we found that the APR-1 dependent stabilization of MT-cortex interactions negatively regulates the pulling forces acting on the anterior centrosome in wild-type zygotes. Our study identifies APR-1 as an attenuator of spindle-pulling forces, and improves our understanding of how cortical polarity precisely regulates spindle positioning during asymmetric cell division.

## Results and Discussion

### APR-1/APC localizes asymmetrically to the cell cortex in a PAR and Frizzled protein dependent manner

We have previously shown that APR-1 localizes asymmetrically to the anterior cortex in the EMS blastomere at the six-cell stage and in post-embryonic seam cells, in response to Wnt signals that regulate the asymmetry of these divisions (31, 32). While analyzing GFP::APR-1 localization in early embryos, we noticed that APR-1 is also asymmetrically localized in the zygote, called P0, where roles for Wnt signaling have not been reported. APR-1 formed dot-like particles that were enriched within the anterior cortex throughout P0 cell division (APR-1 asymmetry) (Figure 1A). We quantified the number of APR-1 dots by counting the fluorescent foci with a signal above a threshold (see Materials and methods). The foci numbers changed from prophase to metaphase, and from anaphase to telophase. Nevertheless, we observed anterior enrichment of APR-1 foci throughout mitosis (Figure 1A and 1D).

**Figure 1.**
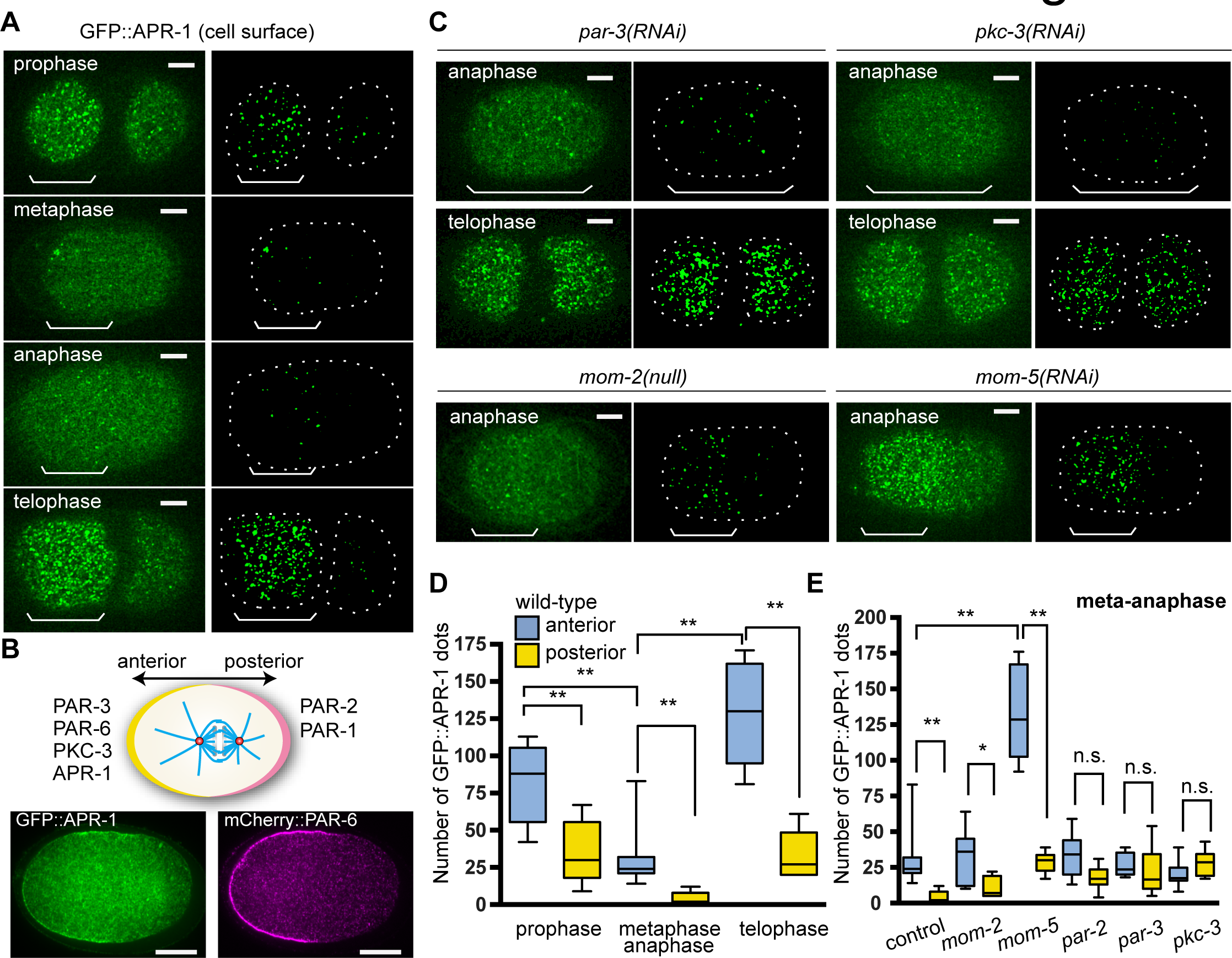
The Par-aPKC system and Frizzled signaling regulate APR-1 asymmetric localization during zygote division. (A) GFP⸬APR-1 signals on the cell surface in different mitotic stages. In the right panels, computationally detected APR-1 dots are shown (see Material and Methods).(B) GFP⸬APR-1 and mCherry⸬PAR-6 localizations in the cell midplane during asymmetric cell division. GFP signal was amplified by the anti-GFP immunostaining. Schematic drawing shows polarized protein localizations. (C) GFP⸬APR-1 signals on the cell surface in *mom-2(null)* mutants and *mom-5, par-2* or *par-3* RNAi embryos. (D) Quantified numbers of GFP⸬APR-1 dots on the anterior and posterior cell cortex of wild-type embryos in different mitotic stages. n = 5, 10, 5 from left to right. (E) Quantified numbers of GFP⸬APR-1 dots at metaphase or anaphase in RNAi embryos. n = 10, 7, 10, 9, 10, 10, from left to right. Ends of whiskers indicate minimum or maximum values. Double asterisk, asterisk and n.s. indicates: p < 0.01, p < 0.05 and p > 0.05 (One-way ANOVA with Holm-Sidak’s multiple comparison test). Scale bars are 10 μm.

It is well-established that the Par-aPKC system generates anterior-posterior (A-P) cell polarity to regulate the asymmetric division of P0, through interactions between anterior (PAR-3, PAR-6, PKC-3) and posterior (PAR-2, PAR-1) *partitioning defective* (PAR) proteins at the cell cortex (Figure 1B; 33). We found that APR-1 asymmetry in P0 was disrupted after RNAi knockdown of *par-3, pkc-3* or *par-2* (Figure 1C, 1E, and Figure S1), suggesting that its asymmetry is established through the Par-aPKC system.

In EMS and seam cells, the establishment of APR-1 asymmetry depends on Wnt proteins (31, 32). In P0, MOM-2 is the only Wnt protein that is maternally provided as mRNA (34), although the mRNA appears not to be translated until the 4cell stage (35). As expected, we found that APR-1 localization was not affected in *mom-2(or309)* null mutants, suggesting that the APR-1 asymmetry in P0 does not require Wnt ligands (Figure 1C, 1E, and Figure S1).

Despite the lack of a requirement for MOM-2/Wnt, we observed altered APR-1 localization after RNAi knockdown of downstream Wnt signaling components. Specifically, knockdown of the Frizzled receptor MOM-5 or simultaneous inhibition of the Dishevelled homologs, DSH-2 and MIG-5, increased the numbers of APR-1 foci at metaphase/anaphase in both the anterior and posterior cortex without altering APR-1 expression levels (Figure 1C, 1E, Figure S1, and Figure S2A). Inhibition of WRM-1/β-catenin did not affect APR-1 localization, and *mom-5(RNAi)* as well as *dsh-2;mig-5(RNAi)* embryos still showed APR-1 asymmetry (Figure 1C, 1E, and Figure S1). DSH-2 localizes to the posterior cell cortex during Wnt-dependent asymmetric cell divisions later in development (31, 36). In contrast, DSH-2 localization in P0 was not asymmetric (Figure S2B), consistent with the lack of Dishevelled requirement in APR-1 asymmetry. Interestingly, inhibition of the Axin homolog PRY-1 and Casein kinase homolog KIN-19 resulted in loss of APR-1 asymmetry only during meta-anaphase, suggesting their partial requirement in the establishment or maintenance of APR-1 asymmetry (Figure S1B and S1C).These results are consistent with observations at a later developmental stage (37). We conclude that APR-1 asymmetry in P0 is established by the Par-aPKC system with partial involvement of Axin and Casein kinase, while Frizzled and Dishevelled negatively regulate the levels of cortical APR-1.

### APR-1 asymmetrically suppresses centrosome movements during P0 cell division

The Par-aPKC system independently regulates two P0 asymmetries: the segregation of cell fate determinants (e.g. PIE-1 and PGL-1) and posterior mitotic spindle displacement and thereby asymmetric cell cleavage. In *apr-1(RNAi)* embryos, GFP::PIE-1 segregated into the posterior daughter cell as in wild-type embryos, indicating that APR-1 is not involved in cytoplasmic determinant localization (Figure S2C). In contrast, *apr-1(RNAi)* embryos showed abnormal spindle oscillations. In wild type P0, posterior spindle displacement (represented by centrosome movements along the A-P axis) starts during metaphase and continues during anaphase when it coincides with transverse oscillations (represented by centrosome movements along the transverse axis) of the two spindle poles (Figure 2A, 2B, 2D, 2E). The posterior spindle pole oscillates more vigorously than the anterior pole (Figure 2B, 2E and Video 1), as a result of higher posterior than anterior cortical pulling forces (38). In *apr-1(RNAi)* embryos, spindle movements were exaggerated: in some embryos, the mitotic spindle moved back and forth along the A-P axis (Figure 2C, 2D, and Video 2), and in some cases, the anterior spindle pole exhibited excessive transverse oscillations, visible by the increased frequency and amplitude of the spindle pole tracks (Figure 2C, 2E and Video 2). As a result, the total distance traveled by the anterior centrosome significantly increased compared to that in control embryos (Figure 2F). These data indicate that APR-1 suppresses anterior spindle pole movements and thereby control spindle positioning during anaphase.

**Figure 2.**
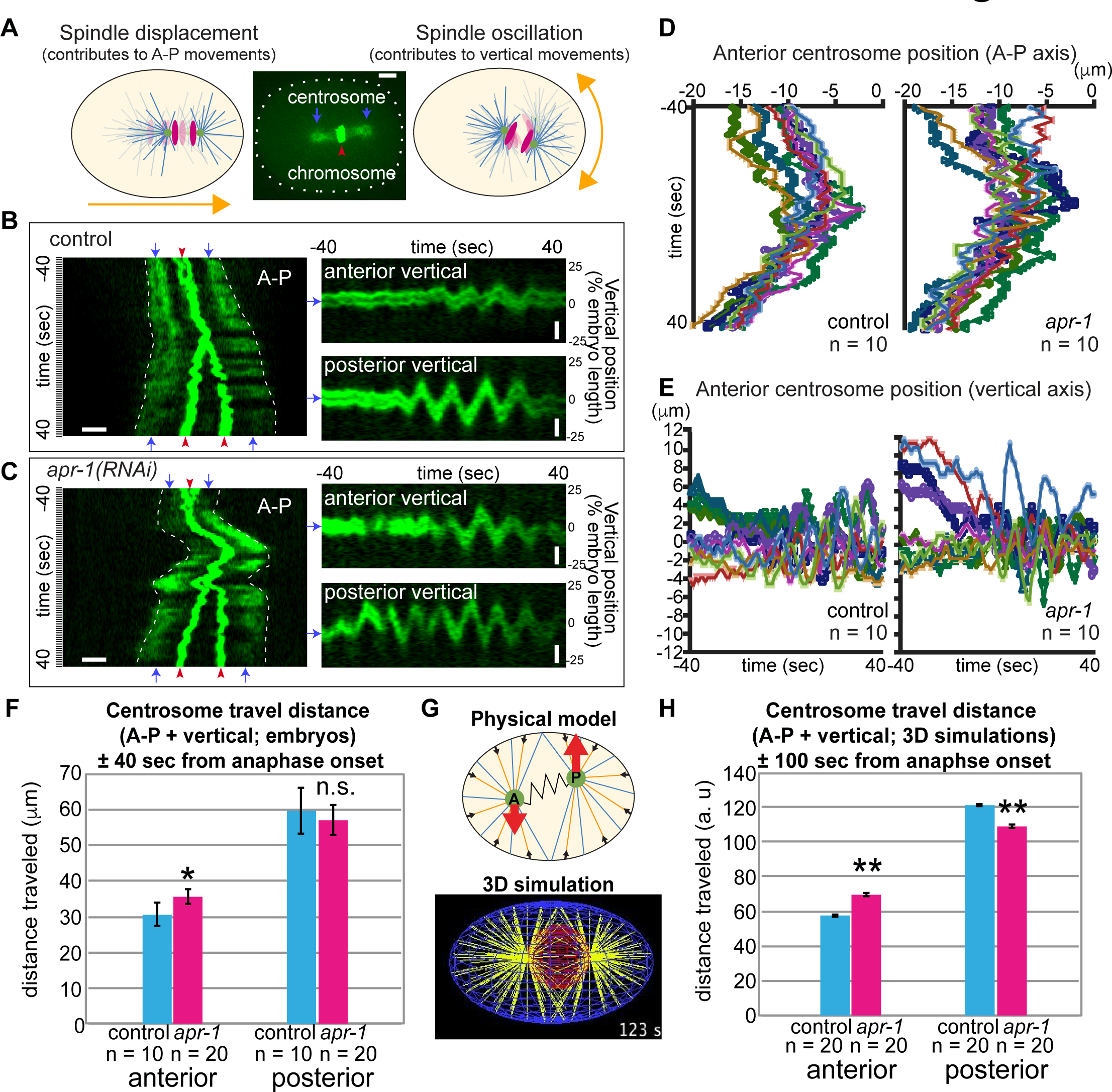
APR-1 asymmetrically suppresses centrosome movements during the P0 cell division. (A) Schematic drawings of spindle movements along the A-P and transverse axes. Spindle displacement and oscillations contribute mainly to the movements along the A-P and transverse axes, respectively. Blue arrows and red arrowhead indicate centrosomes (gamma-tubulin) and chromosomes (Histone H2B), respectively. (B, C) Centrosome movements in A-P (left panels) and transverse (right panels) axes in control (B) and *apr-l(RNAi)* ± 40 second around anaphase onset (C). Kymographs (stack of line images of each time point) were made to show centrosome movements along the A-P and transverse axes separately. (D, E) Anterior centrosome position during cell division along the A-P (D) and vertical axes (E). Cell centers are position zero. (F, H) Total distances for movements of the anterior and posterior poles in living embryos (F) and in 3D simulations (H). (G) Physical model used for 3D simulations. A and P indicate the anterior and posterior spindle poles harboring shrinking MTs (orange) and elongating MTs (blue). Red and black arrows indicate centrosome movements and cortical force generation. For each MT catastrophe at the cortex, the average pulling forces acting on a single MT at the posterior are stronger than those at the anterior, due to the different probabilities of MT-force generator interactions (see Materials and methods). Error bars show 95% CI. Double asterisk and n.s. indicates: p < 0.01 and p > 0.05 compared to control (Kruskal-Wallis test followed by Dunn’s multiple comparison test). Scale bars indicate 5 μm.

In *mom-5(ne12)* null mutant embryos, in which APR-1 levels were increased at both the anterior and posterior cortex, we observed reduced posterior spindle pole oscillations (Figure S3A and S3B). However, spindle pole oscillations were not restored in *apr-1(RNAi); mom-5(null)* embryos (Figure S3B). These results suggest APR-1-independent functions of MOM-5 that influence spindle movements. Because of this, we could not determine the effects of excess cortical APR-1 on spindle pole movements in the *mom-5(null)* background. However, in other aspects of spindle dynamics described below, elevated cortical APR-1 localization potentiated APR-1 function.

### APR-1 asymmetrically stabilizes microtubule-cortex interactions

As mammalian APC (39) and *C. elegans* APR-1 in the EMS cell (32) can stabilize MTs, we hypothesized that anteriorly enriched APR-1 in the P0 cell may also increase MT stability at the cell cortex to regulate asymmetric spindle movements. To assess this possibility, we analyzed the MT-cortex interactions using live imaging of GFP⸬β-tubulin expressing embryos. In kymographs of midplane images, astral microtubules appear to persist longer on the anterior cell cortex than on the posterior, consistent with previous observations (Figure 3A; 40). To precisely quantify MT-plus end residence time at the cortex, we measured the duration of GFP⸬β-tubulin foci on the flattened cell surface (Figure 3B). Most of the GFP⸬β-tubulin foci initially co-localized with the EB1-related plus-end binding protein EBP-2 (96.1%; n = 255), confirming that the foci represent MT plus-ends. Shortly after the cortical attachment, EB1 dissociates from MT plus-ends, while some MTs remained at the cortex after the release of EB1 (Fig. 3B and 3D). The numbers of such long-lived microtubule plus-ends were higher anteriorly, accounting for the asymmetry in cortical MT residence time in wild-type zygotes (Figure 3B-3D; red arrows in 3C, Video 3 and Video 4).

**Figure 3.**
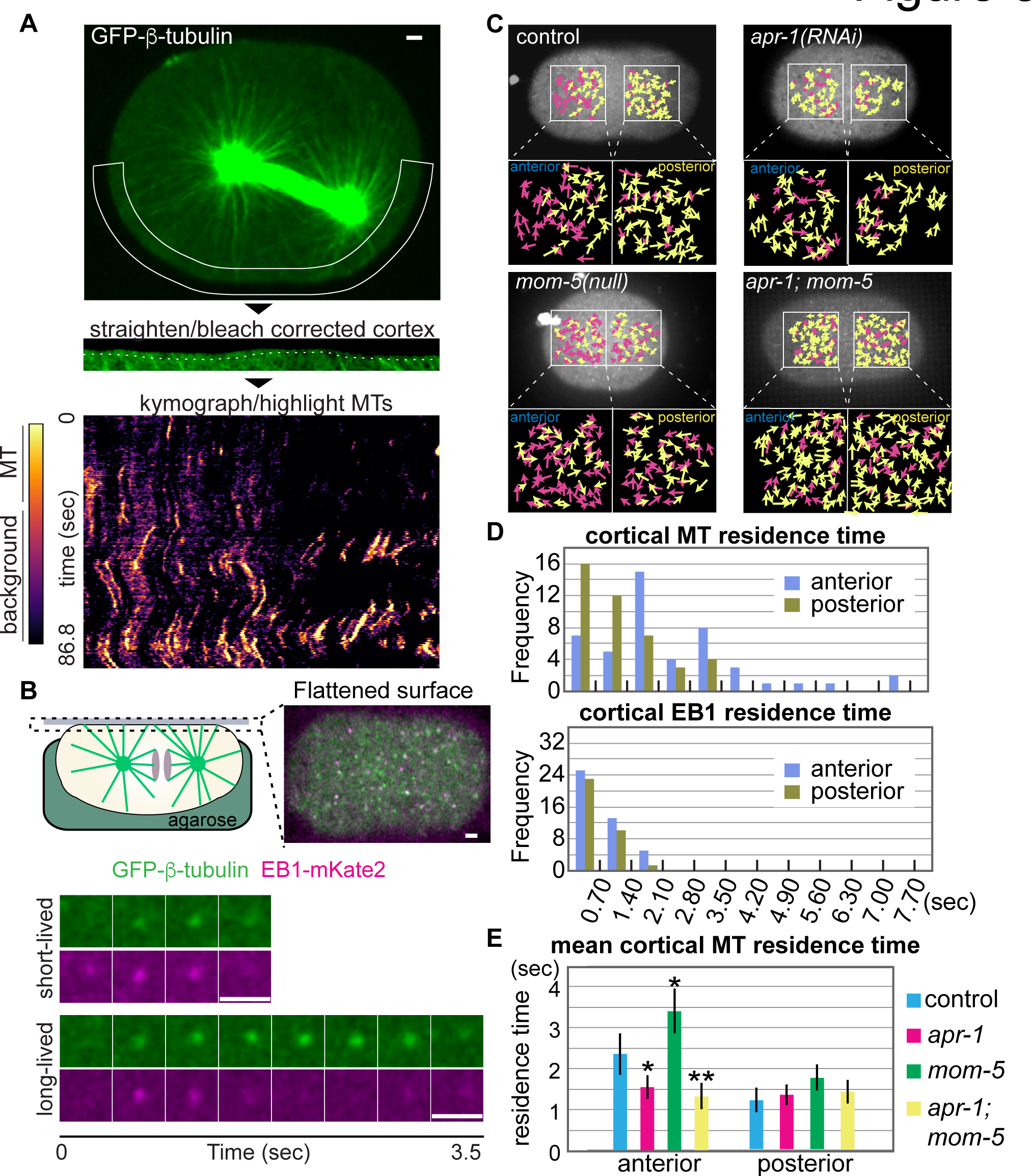
APR-1 asymmetrically stabilizes microtubule-cortex interactions. (A) Cortical MT dynamics. Cortical area outlined by the solid line in top figure was extracted, straightened, and corrected for photobleaching. This cortical area, depicted by the dotted line (middle), was used to generate a kymograph (bottom). Color code of the kymograph was changed to highlight MTs. (B) Measurement of cortical MT residence. The embryos were mounted on agarose pads and flattened by coverslips to visualize cortical microtubule ends in a single focal plane. Examples of short and long-lived foci were shown below with simultaneous imaging of GFP⸬β-tubulin and EB1⸬mKate2. (C) Cortical microtubule dots in the indicated genotypes during metaphase-anaphase. Images are max projection of cortical GFP⸬β-tubulin for 60 frames (42 sec). Yellow and Magenta arrows indicate the MT dots whose residence time was shorter and longer than 2.1 sec, respectively. See also Video 3, 5-7. (D) Distribution of quantified cortical MT or EB1 residence time in wild-type animals. (E) Mean cortical MT residence time of indicated genotypes. n = 47, 42, 77, 67, 64, 61, 37, 44, from left to right. Error bars show 95% CI. Double asterisk and asterisk indicate: p < 0.01 and p < 0.05 compared to control (Kruskal-Wallis test followed by Dunn’s multiple comparison test). Scale bars indicate 2.5 μm.

Notably, the MT residence time at the anterior cortex was significantly lower in *apr-1(RNAi)* embryos than in the wild type (Figure 3C, 3E and Video 5). In contrast, *mom-5* mutants with excess cortical APR-1 showed an increased MT residence time at both the anterior and posterior cell cortex (Figure 3C, 3E and Video 6). RNAi knockdown of *apr-1* overcame this *mom-5* phenotype, reducing MT cortical residence throughout the cortex (Figure 3C, 3E and Video 7). Thus, APR-1 stabilizes microtubule-cortex interactions and acts downstream of MOM-5 (Figure 4D).

**Figure 4.**
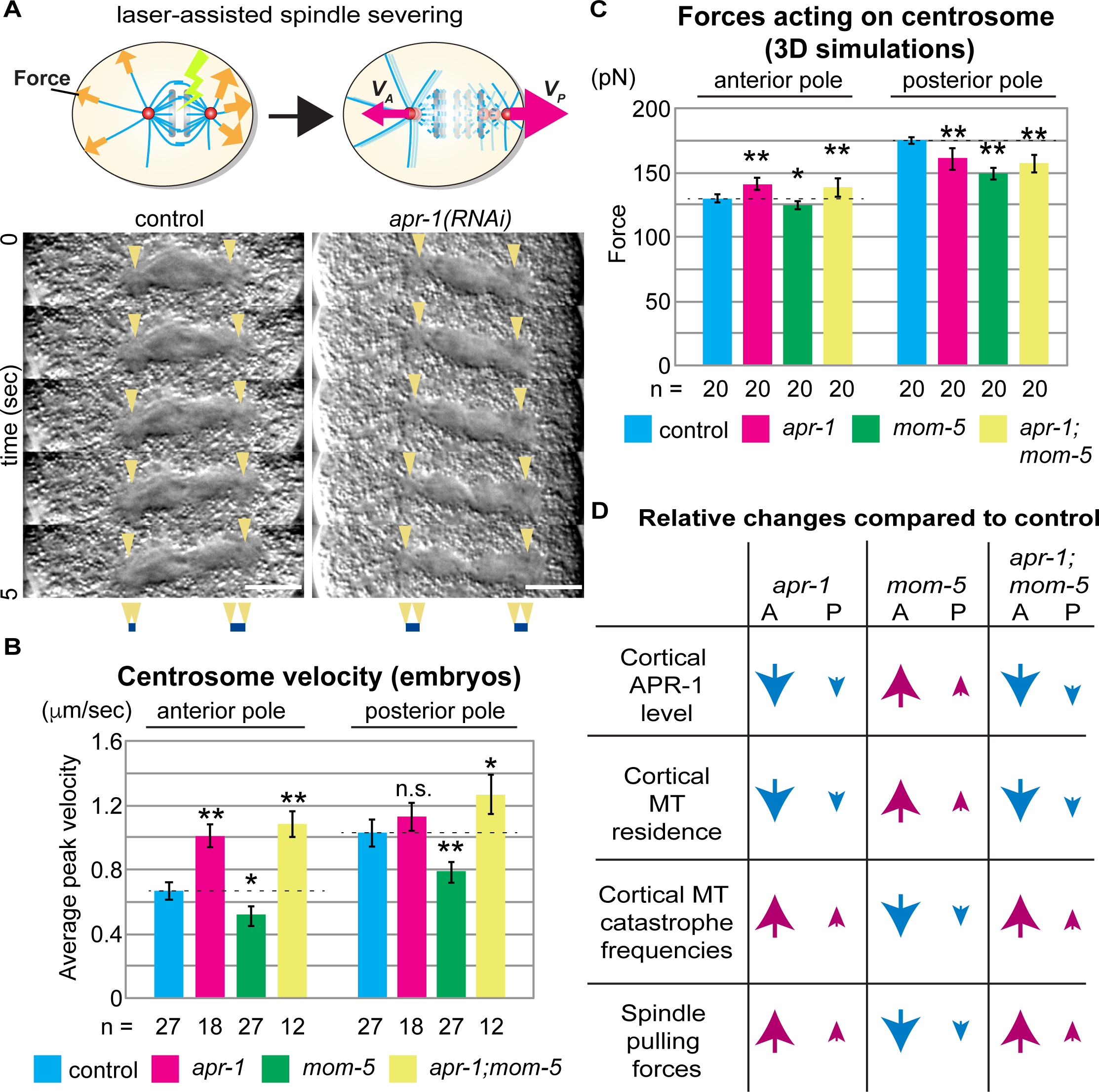
APR-1 asymmetrically attenuates pulling forces acting on the mitotic spindle. (A) Spindle severing experiments. The midzones of mitotic spindles were severed by laser irradiation around anaphase onset (upper left panel). Upon spindle severing, spindle remnants moved at different velocities depending on the net strength of pulling forces (upper right panel). Montages of dissected spindle dynamics were shown in the bottom panels as DIC images; spindle poles devoid of yolk granules were indicated by arrowheads. (B) Average peak velocity of spindle poles after spindle severing. (C) The average of outward pulling forces over 5 sec from anaphase onset (t = 100 s) for 20 independent simulations. Error bars show 95% CI. Double asterisk and asterisk indicate: p < 0.01 and p < 0.05 compared to control (one-way ANOVA with Holm-Sidak’s method). (D) Summary of relationships between cortical APR-1 level, cortical MT residence, cortical MT catastrophe frequencies, and spindle pulling forces. Scale bars indicate 10 μm.

### APR-1 asymmetrically attenuates pulling forces acting on the mitotic spindle

The exaggerated anterior spindle pole movements in *apr-1(RNAi)* embryos implicate APR-1 in spindle-pulling force regulation. We investigated this possibility using spindle severing assays (Figure 4A; 41). After cutting the spindle midzone with a UV laser, the average peak velocities of the anterior and posterior spindle poles moving toward the cell cortex were calculated (Figure 4A). In control embryos, the posterior spindle pole moved faster than the anterior pole, as expected (Figure 4A, 4B, and Video 8). In *apr-1(RNAi)* embryos, we observed an increased average peak velocity specifically for the anterior spindle pole (Figure 4A, 4B, and Video 8). In *mom-5(null)* embryos with excess cortical APR-1, both the anterior and posterior spindle poles showed reduced average peak velocities (Figure 4B and Video 8). Combined *apr-1(RNAi);mom-5(null)* embryos showed increased average peak velocities and resembled *apr-1(RNAi)* embryos (Figure 4B and Video 8). These results indicate that the cortical levels of APR-1 inversely correlate with spindle-pulling forces and suggest a role for APR-1 as cortical pulling force attenuator (Figure 4D).

### APR-1-dependent stabilization of MTs accounts for reduced pulling forces on the anterior spindle pole

We have shown that APR-1 is enriched at the anterior cell cortex, promotes cortical MT residence times anteriorly, and suppresses both spindle-pulling forces and anterior spindle pole oscillations, raising the possibility that these processes are mechanistically linked. It has been shown that cortical pulling forces are generated when MTs reaching the cortex meet dynein and undergo catastrophe (transition from MT plus end growth to rapid shrinkage) (42). Therefore, we hypothesized that cortical APR-1 reduces the MT catastrophe frequency and thereby attenuates force generation and spindle movement. However, it is not clear whether the magnitude of APR-1-dependent cortical MT stabilization is sufficient to suppress spindle movement.

We decided to examine this issue using numerical simulation. First, we estimated MT catastrophe frequencies from their cortical residence time (Supplementary Table 1, Figure S4). In control embryos, the estimated catastrophe frequency at the anterior cortex was about half of that at the posterior cortex. Such a reduced catastrophe frequency was not detected at the anterior cortex of *apr-1(RNAi)* embryos, indicating that in wild type embryos the catastrophe frequency is suppressed by APR-1.

We set the rescue frequency of all MTs high, so that soon after the MTs start to shorten, they regrow to reach the cortex (Supplemental Table 2). This assumption was introduced to make the number of MTs reaching the cortex almost constant regardless of the differences in catastrophe frequencies between anterior and posterior, which is the case in living embryos (Video 3). Without this assumption, the number of MTs reaching the cortex should be ~2-fold higher at the anterior because the catastrophe frequency is about half of the posterior catastrophe frequency. The mechanistic basis of this assumption is provided by the *in vivo* observation that individual microtubules appear to form bundles, and multiple EB1 tracks move along a bundled fiber toward the cell cortex, making rescue frequency of the fiber higher than individual MTs (Video 4), which is consistent to the previous observation (43).

We conducted 3-dimensional simulations of spindle movements. As in previous simulations (44-47), the spindle moves as a result of three kinds of forces acting on astral MTs that radiate from each spindle pole (Figure 2G). First, all MTs generate pulling forces proportional to their length (“cytoplasmic pulling force”). This force is important for positioning of the spindle in the cell center during mitotic prophase (45, 48, 49), and is also critical for oscillation (38). Second, MTs that reach the cell cortex generate the pulling force at their plus ends only when they undergo catastrophe (“cortical pulling force”). The current theory for the basis of oscillation is that when the spindle poles move toward one side, the pulling force from that side becomes stronger (“positive feedback” or “negative friction”), while the opposing centering force also increases (38, 50, 51). With this mechanism, the spindle is not stabilized at the center but oscillates. In our model, the frequency of the force generation depends on the number of active cortical force generators and the MT residence time controlled by APR-1, both of which have A-P asymmetry. The third force connects the anterior and posterior spindle poles. We assumed a spring-like connection between the poles that was weakened after anaphase onset to mimic the spindle elongation.

Numerical simulations were conducted for control, *apr-1(RNAi),* and *mom-5(null)* situations (Figure S5), by setting the catastrophe frequency to values estimated from experimental data (e.g. 0.31 /s for the anterior and 0.72 /s for the posterior, see Supplementary Table 1). The simulation results indicated that the APR-1-dependent stabilization of MTs is sufficient to suppress oscillation of the anterior pole (Figure 2H). In wild-type simulations, the spindle moved toward the posterior and elongated upon anaphase onset (Figure S5A and Video 9). The oscillations perpendicular to the A-P axis were also reproduced for both spindle poles (Figure S5B). In *apr-1(RNAi)* simulations, in which the catastrophe frequency at the anterior cortex was increased, the amplitude of the anterior spindle pole oscillations was increased (Figure 2H,

Figure S5 and Video 9). Furthermore, the average peak velocities of anterior poles in the severing experiments were also consistent with the forces acting on anterior spindle poles in our simulations (Figure 4C). Overall, the numerical simulations supported the hypothesis that the APR-1-dependent stabilization of MTs at the cortex can suppress spindle pole oscillations through the reduction of force generation.

### Anterior APR-1 and LIN-5 phosphorylation together attenuate spindle pulling forces

We investigated the significance of spindle pulling force attenuators in asymmetric cell division. Along with APR-1, we focused on the LIN-5 protein. LIN-5 interacts with cortical GPR-1/2 and dynein in cortical force generation (52). We have previously reported that anteriorly-localized PKC-3/aPKC phosphorylates LIN-5 to attenuate cortical-pulling forces (53). We edited the *lin-5* genomic locus to substitute four aPKC phosphorylated serine residues with alanine by CRISPR/Cas9-mediated homologous recombination *(lin-5 4A* mutation). In spindle severing experiments, combining *apr-1(RNAi)* and the *lin-5 4A* mutation caused significantly enhanced average peak velocities of the anterior poles as compared to *apr-1(RNAi)* embryos (Figure 5A). Compared to *lin-5 4A* embryos, the increase in anterior peak velocity was not significant (p= 0.07; Figure 5A). However, in contrast to the single mutants, the ratio of anterior to posterior centrosome peak velocities in *apr-1(RNAi); lin-5 4A* double mutants was significantly reduced compared to wild-type controls (Figure 5B). These data suggest that the Par-aPKC-dependent asymmetric localization of APR-1 and phosphorylation of LIN-5 together attenuate cortical pulling forces to generate pulling force asymmetry that positions the mitotic spindle (Figure 5C-5E).

**Figure 5.**
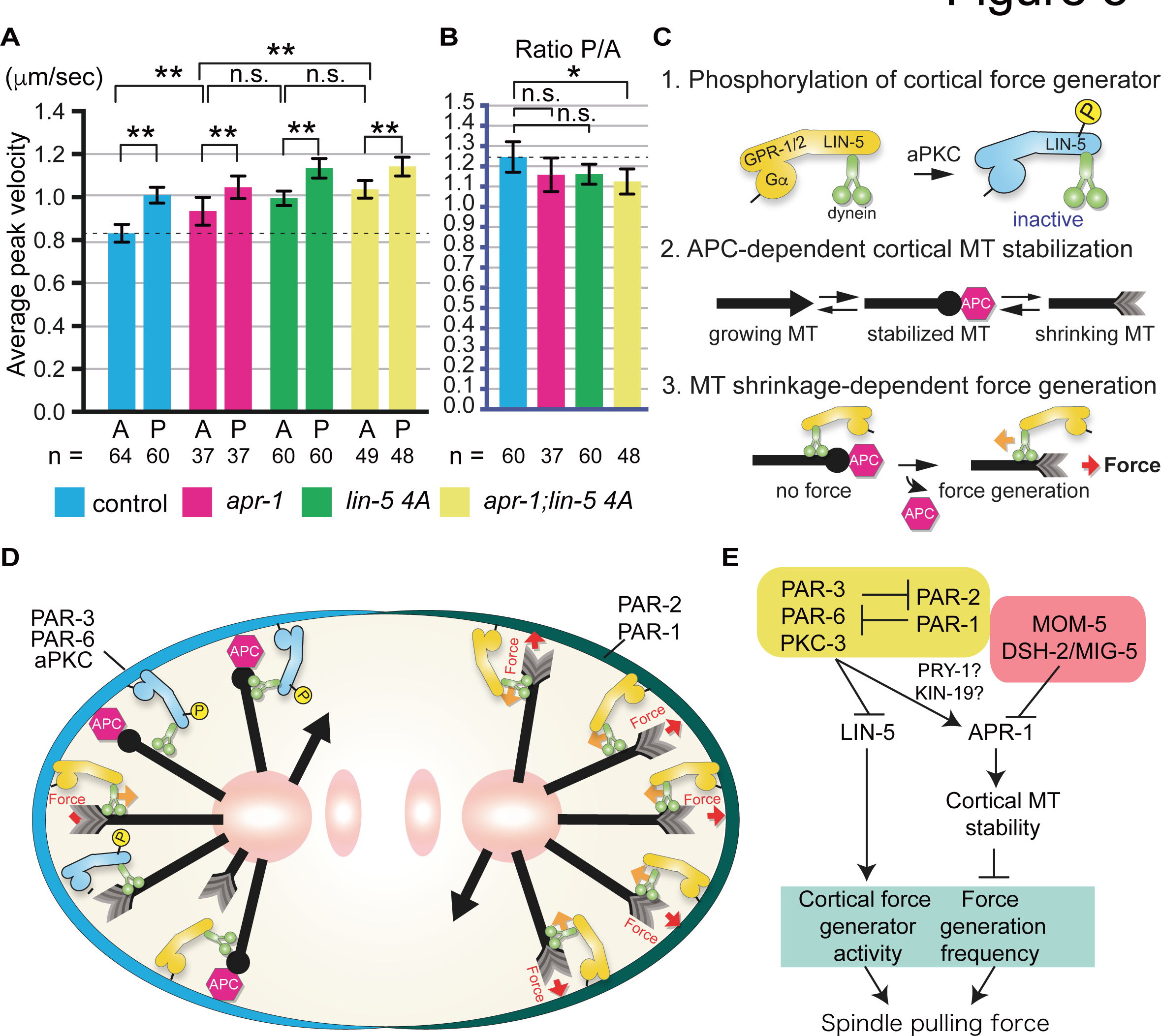
Anterior APR-1 enrichment and LIN-5 phosphorylation together attenuate spindle pulling forces to generate pulling force asymmetry. (A, B) Average peak velocity of spindle poles (A) and their posterior/anterior ratio (B) after spindle severing. Error bars show 95% CI. Double asterisk and asterisk indicate: p < 0.01 and p < 0.05 compared to control (one-way ANOVA with Holm-Sidak’s method). (C) Three elementary processes used in the model described in the panel C. (1) aPKC-dependent LIN-5 phosphorylation results in the inhibition of force generation, (2) Cortical MT stabilization by APC reduces the MT catastrophe frequency and (3) MT shrinkage-dependent force generation is suppressed by step (2).(D) A schematic model of asymmetric spindle force regulation in P0 cell (see text).(E) A diagram of spindle pulling force regulation pathways at the anterior cell cortex.

## Conclusion

In this study, we investigated how the APR-1/APC protein regulates mitotic spindle movements in the *C. elegans* one-cell embryo, a well-established model for asymmetric cell division. We observed that APR-1/APC becomes asymmetrically enriched at the anterior cell cortex, dependent on the Par-PKC-3 polarity pathway. We found that APR-1 attenuates spindle pulling forces, most likely though stabilization of MTs at the anterior cell cortex. In concert, Wnt signaling components MOM-5/Frizzled and Disheveled proteins suppressed cortical accumulation of APR-1, thereby contributing to the correct levels of pulling forces. To test these assumptions, we performed numerical simulations, which closely mimicked the spindle movements in wild-type and mutant embryos. These combined data strongly support the conclusion that MT stabilization by APR-1 contributes to correct spindle positioning. Finally, we provide evidence to suggest that asymmetric APR-1 enrichment and PKC-3 phosphorylation of LIN-5 act in parallel to regulate asymmetric cell division. These conclusions are likely to apply broadly and improve our understanding of the microtubule-associated functions of APC.

Although APC is a component of Wnt signaling, its localization has been reported to be regulated by the Par-aPKC polarity pathway in migrating mammalian astrocytes (54), and during axonal differentiation of developing hippocampal neurons(55), as we observed in the *C. elegans* one-cell embryo. Scratching of astrocyte monolayers in wound-healing assays triggers APC localization to the cell cortex at the leading edge, in response to CDC42-induced Par-aPKC polarity and Wnt5a signaling(56). Interestingly, polarity establishment in this system is followed by centrosome re-orientation through APC-microtubule interactions (54). Thus, the mechanisms that control centrosome positioning through interactions between Par polarity, Wnt signaling, and APC may be conserved across species. The dynamic change in cortical APR-1 levels during P0 cell division is intriguing: this may reflect cell cycle dependent activation of the Wnt signaling pathway as reported in fly and mammalian cultured cells (57).

While the roles of cortical APC have been unclear, it was previously proposed that APC stabilizes microtubules through microtubule plus-end binding protein EB1 (54, 58). Consistently, in the *C. elegans* EMS blastomere, cortical APC stabilizes MT ends coated with EB1 (32). However, a few examples including the present study indicate that cortical APC can stabilize microtubules independently of EB1. First, truncated mammalian APC that lacks the EB1 interaction domain has been shown to localize to the cell cortex and to MTs in epithelial cells (59). In addition, *Drosophila* APC2, which lacks the C-terminal EB1 binding domain, interacts with microtubule plus ends at the cortex and contributes to centrosome segregation (28). In our study, APR-1 at the anterior cortex stabilizes MTs but the mean cortical residence time of EBP-2/EB1 was symmetric. We also observed that the cortical residence time of EB1 is much shorter than that of MTs in P0, as reported previously (43). Therefore, APR-1 at the anterior cortex of P0 likely stabilizes MTs independently of EB1 binding. We observed recently that deleting all EB family members has limited effects on spindle behavior and viability in *C. elegans* (60). Therefore, the microtubule stabilizing effects of cortical APC probably do not depend on EB1 protein interactions.

Mitotic spindle positioning is tightly controlled during embryogenesis, in various adult stem cell divisions, and in symmetric divisions (1, 3, 61). While many studies have focused on the localization of cortical force generators that pull on microtubule plus ends, attenuators of spindle pulling forces may be just as important in creating asymmetry. In fact, a variety of molecular mechanisms appear to suppress spindle pulling forces in the one-cell embryo, including PKC-3-mediated LIN-5 phosphorylation (53), cortical actin (62), and posterior-lateral LET-99 localization (63). This study provides insight into and a physical basis of spindle pulling force attenuation: we found that APC acts as an attenuator of spindle pulling forces, through stabilization of microtubule plus ends at the cortex. Importantly, a similar force attenuator function of APC is potentially used in oriented divisions of *Drosophila* germline stem cells (11), as well as mouse embryonic stem cells (ES cells) attached to Wnt-immobilized beads (64), as these systems exhibit asymmetric APC localizations similar to what we have observed in the *C. elegans* zygote. Our study also implies that not only APC but also other proteins involved in MT stabilization are potential cortical spindle pulling force attenuators.

The observed pulling force attenuation function may be relevant for the chromosomal instability (CIN) phenotype associated with APC loss in human colon cancer (18, 20). Initial studies of cultured mammalian cells associated APC loss and CIN with defective kinetochore-microtubule attachments, although abnormal spindle structures were also observed in APC defective cells (18, 20). In *Drosophila* embryos, APC2 was found to localize predominantly to the cell cortex (65). Chromosome missegregation associated with APC2 loss in such embryos was linked to a cytoskeletal function of APC in centrosome segregation (28). In our study, we found that *C. elegans* APC localizes to the cell cortex where it negatively regulates spindle pulling forces. Consequently, the absence of APC results in increased pulling forces exerted on the spindle poles. Interestingly, defective kinetochore attachments have been shown to cause chromosome segregation defects in *C. elegans,* in a manner dependent on cortical pulling forces (66). Thus, combining these data with our results raises a new and testable hypothesis that increased cortical-pulling forces and abnormal MT-kinetochore interactions synergistically elevate the risk of CIN in developing tumors with APC mutations.

## Materials and methods

### *C. elegans* culture and strains

All strains used in this study were cultured by standard methods (67). Most worms were grown at 20 °C or 22.5 °C and then incubated at 25 °C overnight before the analysis. Worms used for anti-DSH-2 staining were grown at 22.5 °C. Worms carrying PIE-1: : GFP were grown at 15°C and incubated at 25°C overnight before the analysis. The following alleles were used: *mom-2(or309), mom-5(ne12), par-2(it51).* We used *mom-5(ne12)* null mutants for all *mom-5* experiments except those in Figure 1. The following integrated transgenic lines were used: *osIs15* (32) for GFP⸬APR-1; *ruIs32* (68) for GFP⸬H2B; *ojIs1* (69) for GFP⸬β-tubulin; *axIs1462* (70) for GFP⸬PIE-1; *axIs1720* (70) for GFP⸬PGL-1; *tjIs8* for GFP⸬EBP-1; *ruIs57* for GFP⸬tubulin; *ax1928* for mCherry⸬PAR-6 (71);. We also generated EBP-2⸬mKate2 fusion strain ebp-2(or1954[ebp-2⸬mKate2]) and *lin-5 (he260[S729A,S734A,S737A,S739A])strain* by CRISPR/Cas9 genome editing as described below.

### Generation of CRISPR repair templates

For the generation of the *ebp-2*⸬mKate2 strain, CRISPR repair constructs containing 700 bp homologous arms were synthesized as gBlock fragments (Integrated DNA Technologies, Coralville, IA) and assembled into pJET2.1 vector using in-house Gibson Assembly reaction mix (72). For the generation of the *lin-5 4A* strain, CRISPR repair constructs were inserted into the pBSK vector using Gibson Assembly (New England Biolabs, Ipswich, MA). Homologous arms of at least 1500 bp upstream and downstream of the CRISPR/Cas9 cleavage site were amplified from cosmid C03G3 using KOD Polymerase (Novagen (Merck) Darmstadt, Germany). Linkers containing the point mutations were synthesized (Integrated DNA technologies, Coraville, IA). Mismatches were introduced in the sgRNA target site to prevent cleavage of the repair template and knock-in alleles. All plasmids and primers used for this study are available upon request.

### CRISPR/ Cas9 genome editing

Young adults were injected with solutions containing the following injection mix. For *ebp-2*⸬mKate2, 10 ng/μ! pDD162 Peft-3⸬Cas9 with sgRNA targeting C-terminus of *ebp-2* locus (Addgene 47549; 73), 10 ng/μl repair template, and 65 ng/μl selection marker pRF4 were used. For *lin-5* 4A, 50 ng/μl Peft-3⸬Cas9 (Addgene 46168; 74), 50 ng/μl of two PU6⸬sgRNAs targeting the region of the four serine residues to be mutated to alanine, 50 ng/μl repair template and 2.5 ng/μl selection marker *Pmyo-2*⸬tdTomato were used. Progeny of animals that carry selection markers were transferred to new plates 3–4 days post injection. For *ebp-2*⸬mKate2, GFP positive animals were crossed with a strain carrying GFP⸬ tubulin to obtain ebp-2⸬mKate2 with GFP⸬tubulin (EU3068; *ebp-2(or1954[ebp-2⸬mKate2]* II). For *lin-5* 4A, PCRs with primers diagnostic for recombination products at the endogenous locus were performed on F2-F3 populations, where one primer targeted the altered base pairs in the sgRNA site and point mutations and the other just outside the homology arm. The resulting strain (SV1689; *lin-5* (he260[S729A/S734A/S737A/S739A]) II) was crossed with AZ244 (unc-119(ed3) III; *ruIs57)* to obtain the *lin-5* 4A strain with GFP⸬tubulin (SV1690; *lin-5(he260); ruIs57).*

### RNAi

DNA fragments corresponding to nucleotide 848-1547 of the *apr-1* cDNA were amplified and used for the production of the dsRNA and feeding RNAi. For the experiments in Figure 5, we injected the dsRNA into the gonad and worms were subsequently cultured under feeding RNAi at 25 °C for over 16 hrs before dissecting embryos. For the rest of experiments, after injection of the dsRNA into the gonad, worms were incubated at 25 °C without feeding RNAi for over 30 hrs before dissecting embryos.

### Microscopy and analysis of living embryos

All embryos were dissected in an egg salt buffer from gravid hermaphrodites (75). For live imaging except for the experiments in Figure 5, the embryos were mounted on 4 % agar pads under a coverslip and sealed with petroleum jelly. For most of the experiments embryos were observed at room temperature by a CSU10 spinning-disc confocal system (Yokogawa Electric, Musashino, Japan) mounted on an AxioPlan 2 microscope (Carl Zeiss, Oberkochen, Germany) with a Plan-Apochromat 100X 1.4 NA oil immersion lens. The specimens were illuminated with a diode-pumped solid state 488 nm laser (HPU50100, 20mW; Furukawa Electric, Tokyo, Japan). Images were acquired with an Orca ER12-bit cooled CCD camera (Hamamatsu Photonics, Hamamatsu, Japan), and the acquisition system was controlled by IP lab software (2 X 2 binning; Milwaukee, WI). Acquired images were processed with the Image J (76) (NIH) and Adobe Photoshop (Adobe Systems, San Jose, CA). For the experiments in Figure 3B, images were captured with a confocal unit CSU-W with Borealis (Andor Technology, Belfast, Northern Ireland) and dual EMCCD cameras iXon Ultra 897 (Andor Technology) mounted on an inverted microscope Leica DMi8 (Leica Microsystems, Wetzlar, Germany) controlled by Metamorph (Molecular Devices, Sunnyvale, CA). Spindle severing experiments were performed with a Micropoint system (Photonic instruments, St Charles, IL) equipped with a 2 mW pulsed nitrogen laser (model VL-337; Laser Science Inc., Franklin, MA) exciting Coumarin 440 dye. For the experiments in Figure 5, embryos were mounted on 4 % agarose pad dissolved in egg salts buffer and observed by a Nikon Eclipse Ti microscope with Perfect Focus System (Nikon, Tokyo, Japan) equipped with CSU-X1-A1 spinning disk confocal head (Yokogawa Electric) and S Flour 100X 1.3 NA objectives. The specimens were illuminated with Cobolt Calypso 491 nm laser (Cobolt, Solna, Sweden). Spindle severing experiments were performed with 355 nm Q-switched pulsed lasers (Teem Photonics, Meylan, France) with ILas system (Roper Scientific France, Lisses, France/ PICT-IBiSA, Institut Curie). Temperature was maintained at 25°C by INUBG2E-ZILCS Stage Top Incubator (Tokai Hit, Fujinomiya, Japan) on the motorized stage MS-2000-XYZ with Piezo Top plate (ASI, Eugene, OR). Images were acquired with an Evolve 512 EMCCD camera (Photometrics, Tucson, AZ), and the acquisition system was controlled by MetaMorph (Molecular Devices).

### Immunostaining

For the analysis of GFP⸬APR-1 and mCherry⸬PAR-6 colocalization, we performed the freeze-crack method to permeabilize embryos and fixed them in methanol at −20°C for 5 min followed by acetone at −20°C for 5 min. After washing three times with PBS supplemented with 1% tween-20, the embryos were incubated with rabbit polyclonal anti-GFP antibody (1:1000, invitrogen) overnight. After incubation with goat anti-rabbit Fluorescein (1:1000, Invitrogen), embryos were imaged for Fluorescein and mCherry signal. Embryos were fixed and stained with rabbit anti-DSH-2 antibody as described (77).

### Measurement of embryo volumes

The volumes (V) of embryos were estimated from the measured embryo length (X) and width (Y). When three semi-axes of ellipsoid (embryo) in the x, y and z axes are defined as a, b and c, volume of ellipsoid V = 4/3πabc. With the assumption of equal embryo width in the y and z axes, we estimated a, b and c as 0.5X, 0.5Y and 0.5Y and calculated V.

### Statistical analysis

For multiple comparisons, one-way ANOVA with Holm-Sidak’s method and Kruskal-Wallis test followed by Dunn’s multiple comparison test were performed for the data with normal distribution and skewed distribution (judged by F-test), respectively. No statistical method was used to predetermine sample size. The experiments were not randomized. The investigators were not blinded.

### Quantification of the data from fluorescence images

For the quantification of the number of dots formed by GFP⸬APR-1, 8 bit images were processed with Gaussian blur and segmented with the threshold that covers all the visible dots using Fiji. Then number of segments were counted by the Image J plug-in Analyze Particles. For the quantification of total APR-1 level in Figure S2A, 4 successive focal planes including cell center and cell surfaces (corresponding to the upper half of the cell) were combined by the sum projection and average signal intensity of cell region was subtracted by that in the area devoid of embryos. For the generation of kymographs that show the centrosome movements along the anterior posterior axis, (Figure 2B and 2C, left panels), we drew lines passing through both centrosomes (some centers are missing due to the transverse movements) and generated kymographs using Image J function Multi Kymograph. For the generation of kymographs that show centrosome movements along the transverse axis (Figure 2B and 2C, right panels), we first adjusted the center of the centrosome manually and drew a line that passes through the center of the anterior or posterior centrosome and performed the same procedures. Note that kymographs are composed of linear pixels of each frame for all time points that together show the centrosome trajectory over time. For the quantification of spindle movements, the coordinates of the center of the centrosomes were analyzed with the Image J plug-in Manual Tracking. For the generation of kymographs of cortical microtubules, (Figure 3A), we extracted and straightened cortical regions and performed photo-bleach corrections (exponential fit method) by Image-J. The image color map was changed to mpi-inferno with Image-J. For the quantification of cortical residence times of GFP⸬EB1 and GFP⸬β-tubulin, the number of frames from appearance to disappearance of each dot were counted manually. Note that some MT dots whose start and end of cortical localization were unclear were not counted. The average peak velocity after spindle severing was calculated from the distance traveled by the centrosome center.

### 3-dimensional simulation of spindle movement

#### Overview

The simulations included 2 spindle poles connected by a spring with dynamic astral MTs inside a cell. The cell was simulated as an oval with a long axis of 50 μm and two short axes of 30 μm. The initial position of the spindle poles was set in the center of the cell and aligned along the long axis with the distance of 10μm, which corresponds to the size of the spindle. The MTs grow and shrink from the spindle poles stochastically according to the dynamic instability. Depending on the length and configuration of the MTs, 3 kinds of forces act on spindle poles to move them as explained below. From an initial configuration, the configuration of the MTs and the spindle poles was calculated at successive time steps as conducted in previous simulations (44-47). The parameters used are listed in Table S2.

#### Force 1, cytoplasmic pulling forces

All MTs generate pulling force proportional to their length. This force is important to bring the spindle at the cell center (45, 48, 49), and is also critical for oscillation (38). The cytoplasmic pulling force generated for an *i*-th MT was modeled as *F_cytoplasm_*(*i*) = *D* × *L*(*i*) × *F_FG_*(*i*), where *D* is the density of active force generators in the cytoplasm and *L*(*i*) is the length of the MT. *F_FG_*(*i*) is same as in the cortical pulling force. The direction of the force is the same as the direction of the MT. We note that the centering force required for oscillation can also be provided by a force that microtubules produce when they push against the cortex (78) instead of the cytoplasmic pulling force. The detailed mechanisms (i.e. pulling or pushing) of the centering force do not affect the overall behavior of our model.

#### Force 2, cortical pulling forces

MTs that reached the cell cortex generate pulling forces toward their direction only when they start to shrink. The cortical pulling force generated for an *i*-th MT was modeled as *F_cortex_*(*i*) = *N_potential_*(*i*) × *P_active_*(*i*) × *F_FG_*(*i*). Npotential is the number of force generators that can potentially interact with the MT. We set this value at 30 for the posterior cortex and 15 for the anterior cortex. The experimental value of this parameter has not been investigated, but this number is consistent with a previous study estimating that the total number of force generators is less than 50 and the density is double at the posterior cortex compared to anterior cortex (79). *P_active_* is the probability that the potentially interacting force generators are active. A critical assumption to generate robust oscillation here is to model this value high when the spindle pole is approaching the site of the force generator, and low when the spindle pole is leaving (38, 50). In the previous study (38), *P_active_* was defined as *P_active_* = *p_mean_* + (*f*’/*f_c_*) ×*p_mean_*×(1-*p_mean_*)×*v* − τ × (*f*’/*f_c_*) ×*p_mean_* ×(1-*p_mean_*) ×*a*. For simplicity, we neglected the acceleration term (*a*) and fixed the *p_mean_* parameter to 0.5 to see the extensive oscillation (38). We set *f’/f_c_* = 4.0/ *V_max_*, and thus used *P_active_* =0.5 + *v/V_max_*. Here *v* is the velocity of the spindle pole toward the dire4ction of the force generator on the cortex. When *v*<0 We set *P_active_* =0. *F_FG_* is formulated as *F_FG_* = *F_stall_* (1-*v/V_max_*) (38, 45). When *v*>*V_max_*, we set *F_FG_* = 0. In the simulation, force generation for shrinking MTs lasts for 100 steps (1 s).

#### Force 3, forces connecting the two poles

To connect the anterior and posterior spindle poles, which is done by spindle MTs *in vivo*, we treated the spindle as a Hookean spring. The natural length increases proportionally from 10 μm at time zero to 12 μm at *t* = 100 s, which is the onset of anaphase in the simulation. After the onset of anaphase, the natural length increases proportionally to 22 μm at *t* = 200 s. The spring constant is high (1 pN/μm) so that the length of spindle is almost maintained to the natural length.

**Table 1.**
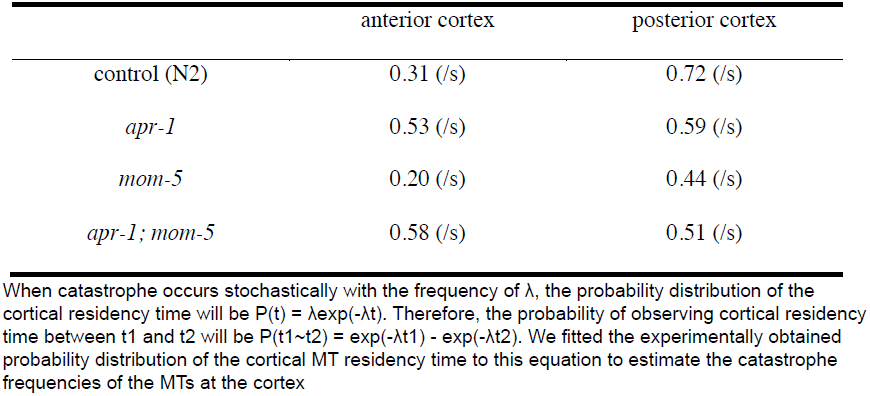
Estimated catastrophe frequencies of the microtubules at the cortex

|  | anterior cortex | posterior cortex |
| --- | --- | --- |
| control (N2) | 0.31 (/s) | 0.72 (/s) |
| *apr-1* | 0.53 (/s) | 0.59 (/s) |
| *mom-5* | 0.20 (/s) | 0.44 (/s) |
| *apr-1; mom-5* | 0.58 (/s) | 0.51 (/s) |

When catastrophe occurs stochastically with the frequency of $\lambda$ , the probability distribution of the cortical residency time will be $P(t) = \lambda \exp(-\lambda t)$ . Therefore, the probability of observing cortical residency time between $t_1$ and $t_2$ will be $P(t_1 \sim t_2) = \exp(-\lambda t_1) - \exp(-\lambda t_2)$ . We fitted the experimentally obtained probability distribution of the cortical MT residency time to this equation to estimate the catastrophe frequencies of the MTs at the cortex

**Table 2.**
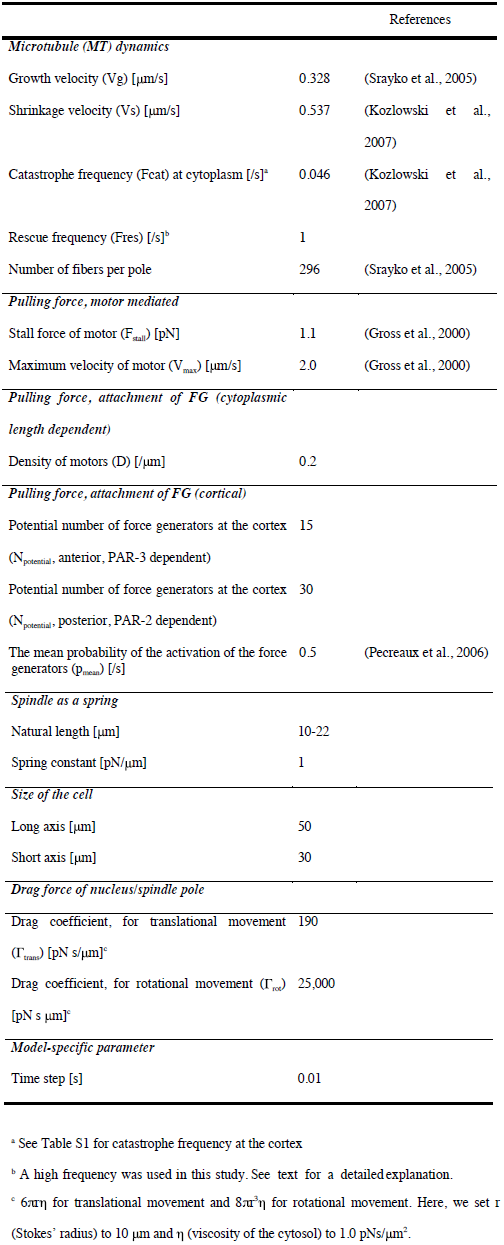
Parameter values used in the simulation

## Acknowledgements

We thank Nancy Hawkins for the anti-DSH-2 antibody, the ***Caenorhabditis*** Genetics Center (funded by the NIH Office of Research Infrastructure Programs; P40 OD010440) for strains. This work was supported by the Netherlands Organization for Scientific Research (NWO) research program 821.02.001 to SvdH, NIH grant R01GM049869 to B.B., by the Human Frontier Science Program and NIG-JOINT (2013-A60) to K.S., by the Uehara Memorial Foundation to H.S., and Grants-in-Aid for Scientific Research from the Ministry of Education, Culture, Sports, Science, and Technology of Japan to H.S (JP22127005) and A.K. (JP15H04732 and JP15KT0083).

---

**Figure S1.**
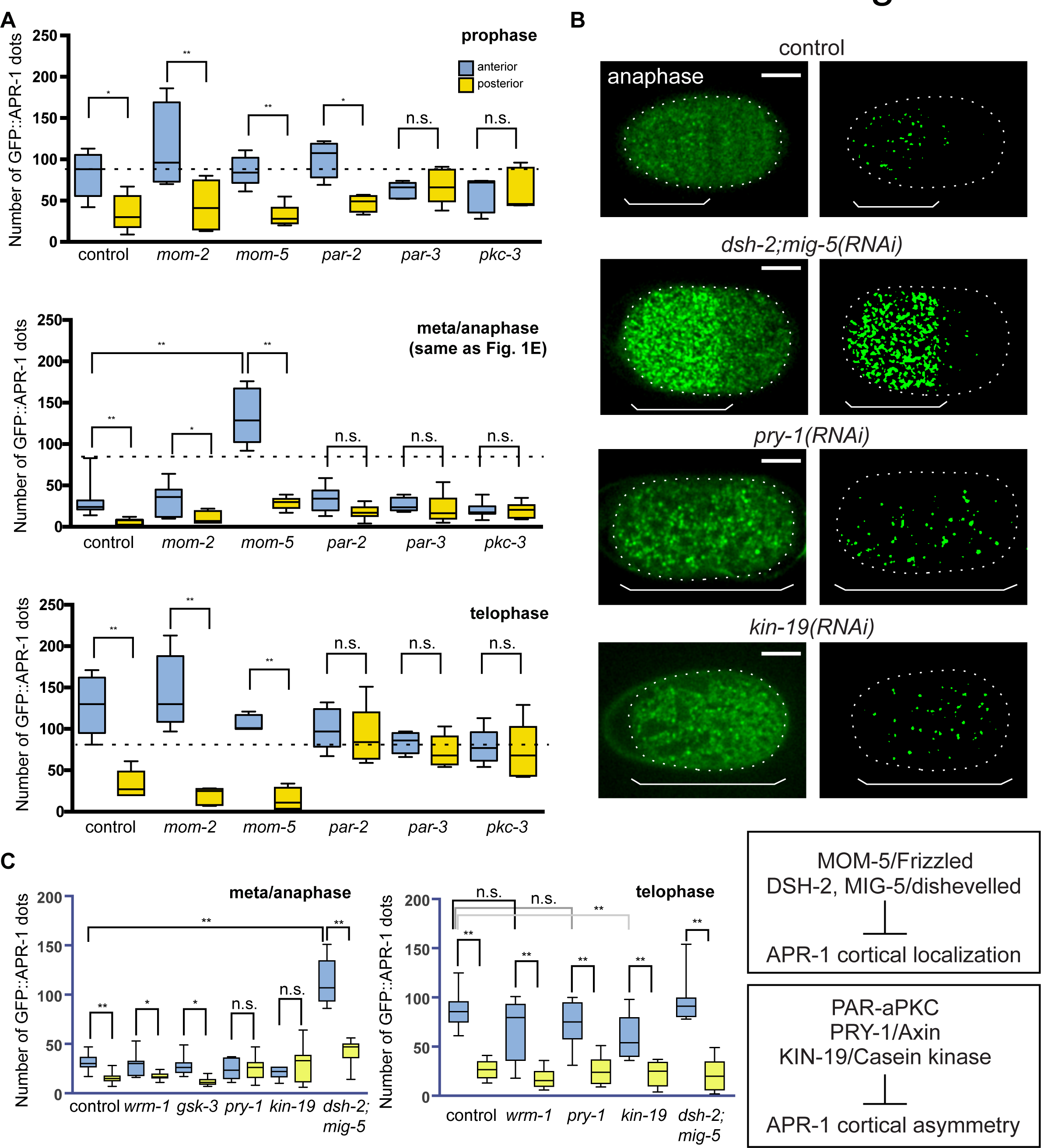
Temporal and genetic regulation of cortical GFP⸬APR-1 localization. (**A, C**) Quantified numbers of GFP⸬APR-1 dots on the anterior and posterior cell cortex are shown for prophase, metaphase, anaphase and telophase of control and RNAi embryos. (B) APR-1 dots in the indicated RNAi experiments. Left and right panels are original and computationally segmented binary images, respectively. Ends of whiskers indicate minimum to maximum values. Double asterisk, asterisk and n.s. indicates: p < 0.01, p < 0.05 and p > 0.05 (One-way ANOVA with Holm-Sidak’s multiple comparison test). Scale bars indicate 10 μm.

**Figure S2.**
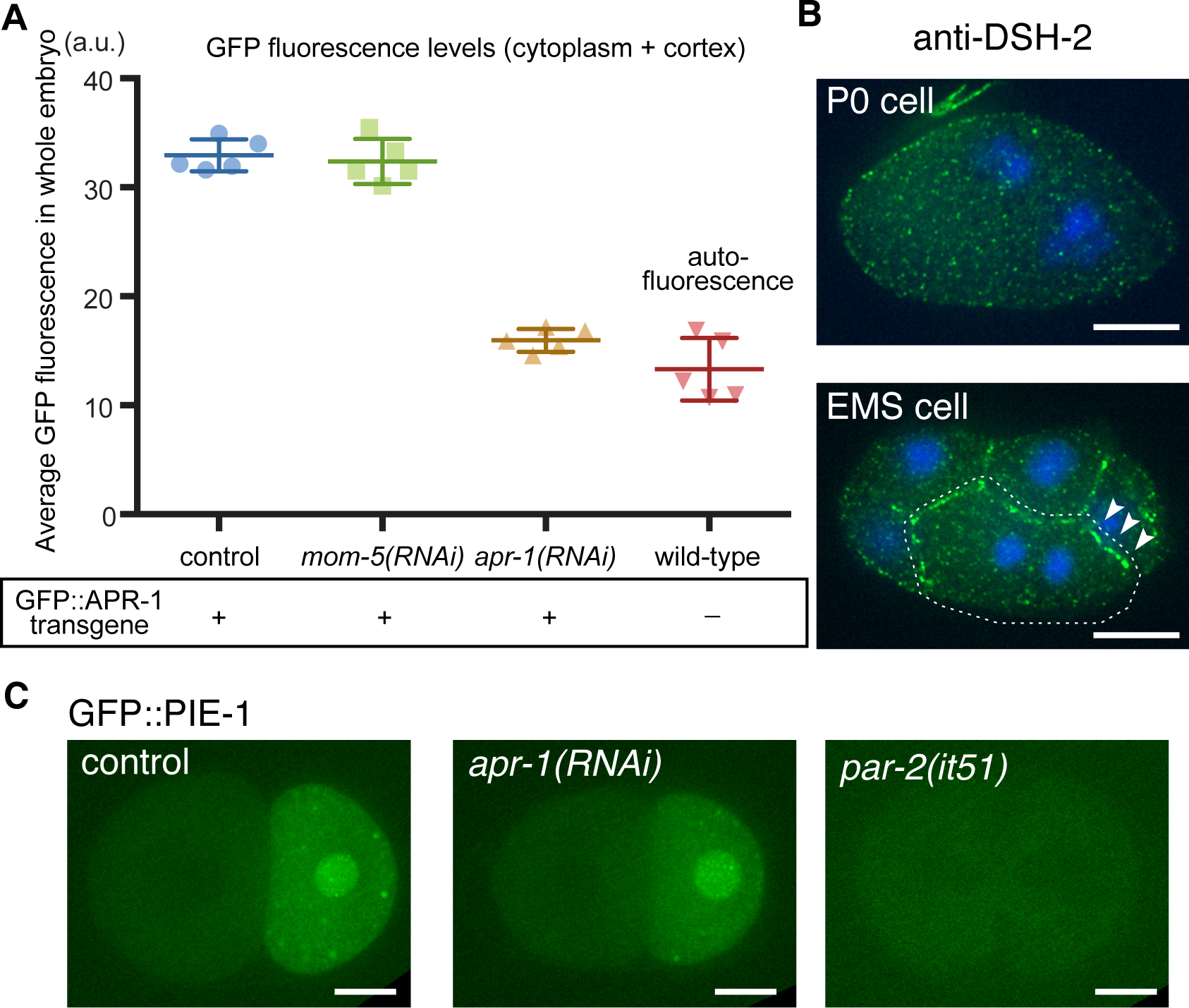
Roles of Wnt signaling in P0 cell division. (A) APR-1 level after RNAi experiments. GFP fluorescence intensity per area of the whole embryo including the cell cortex and cytoplasm were measured and shown. Signal in wild-type indicates autofluorescence. (B) Immunofluorescence images of the DSH-2 protein during P0 and EMS cell division. Blue is DAPI staining. In EMS, the DSH-2 protein is enriched at the cell boundary between EMS and P2 (arrowheads) while no asymmetry was observed in P0. (C) Localizations of the cell fate determinant GFP⸬PIE-1 in the indicated genotypes. Control and *apr-1(RNAi)* shows PIE-1 enrichment in the posterior blastomere P1. In the *par-2* mutant, PIE-1 asymmetry was lost. Scale bars indicate 10 μm.

**Figure S3.**
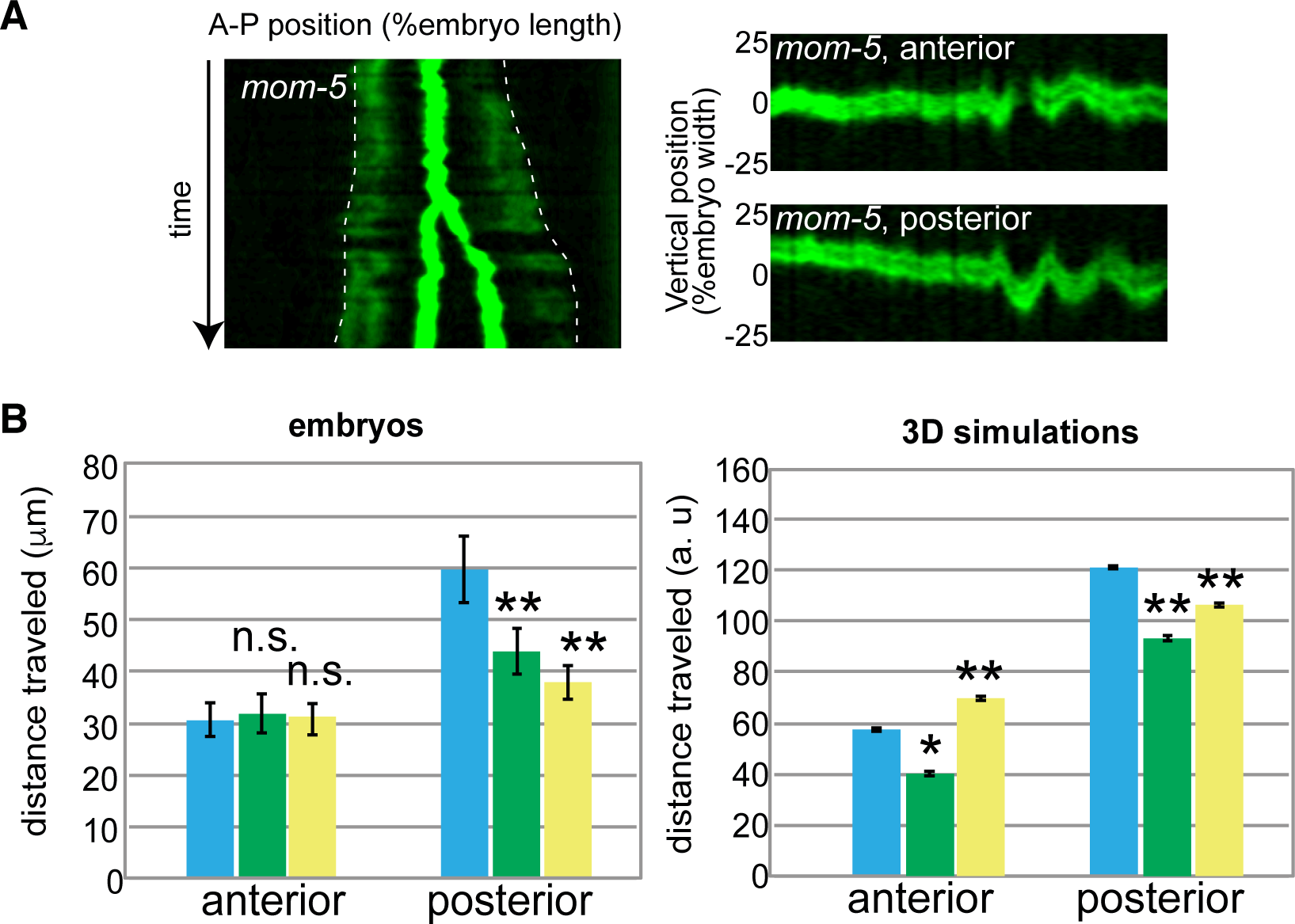
Effects of *mom-5(RNAi)* on spindle pole movements and embryo sizes. (A) Kymographs of the spindle movements in *mom-5(RNAi).* Kymographs were generated as in Figure 2.(B) Distance traveled by the anterior or posterior spindle poles. Total distance centrosome traveled for ± 40 sec and ± 100 sec from anaphase onset were shown for *in vivo* measurements (left) and 3D simulations (right). Error bars show 95% CI. Asterisk indicate p < 0.05 compared to control (Kruskal-Wallis test followed by Dunn’s multiple comparison test).

**Figure S4.**
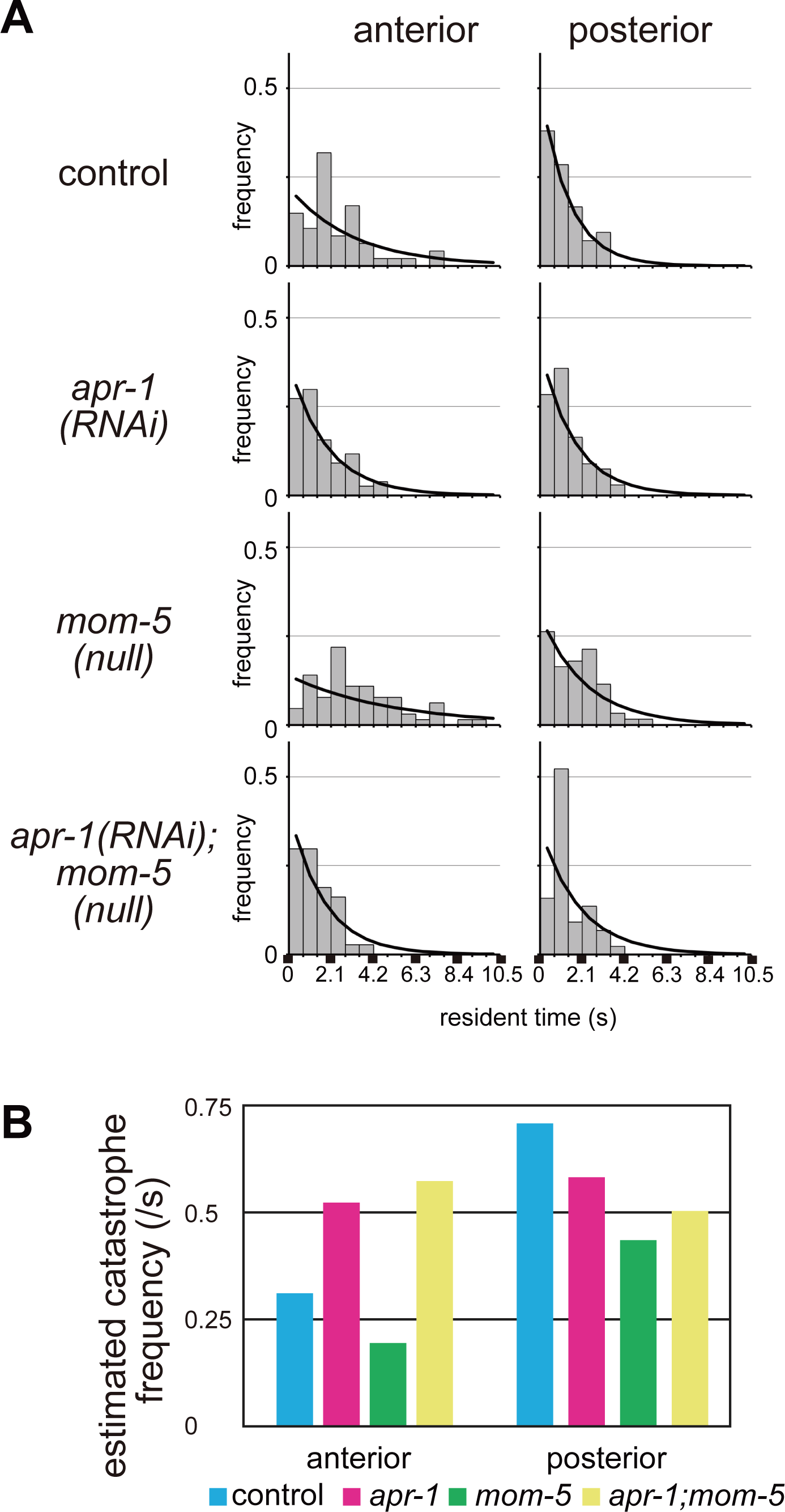
Estimation of microtubule catastrophe frequencies at the cortex. (A) Frequencies of MT residence times at the cell cortex observed experimentally (histograms) and predicted from the estimated catastrophe frequencies (black lines). (B) Estimated catastrophe frequencies for indicated genotypes. The data is the same as in Supplementary Table 1.

**Figure S5.**
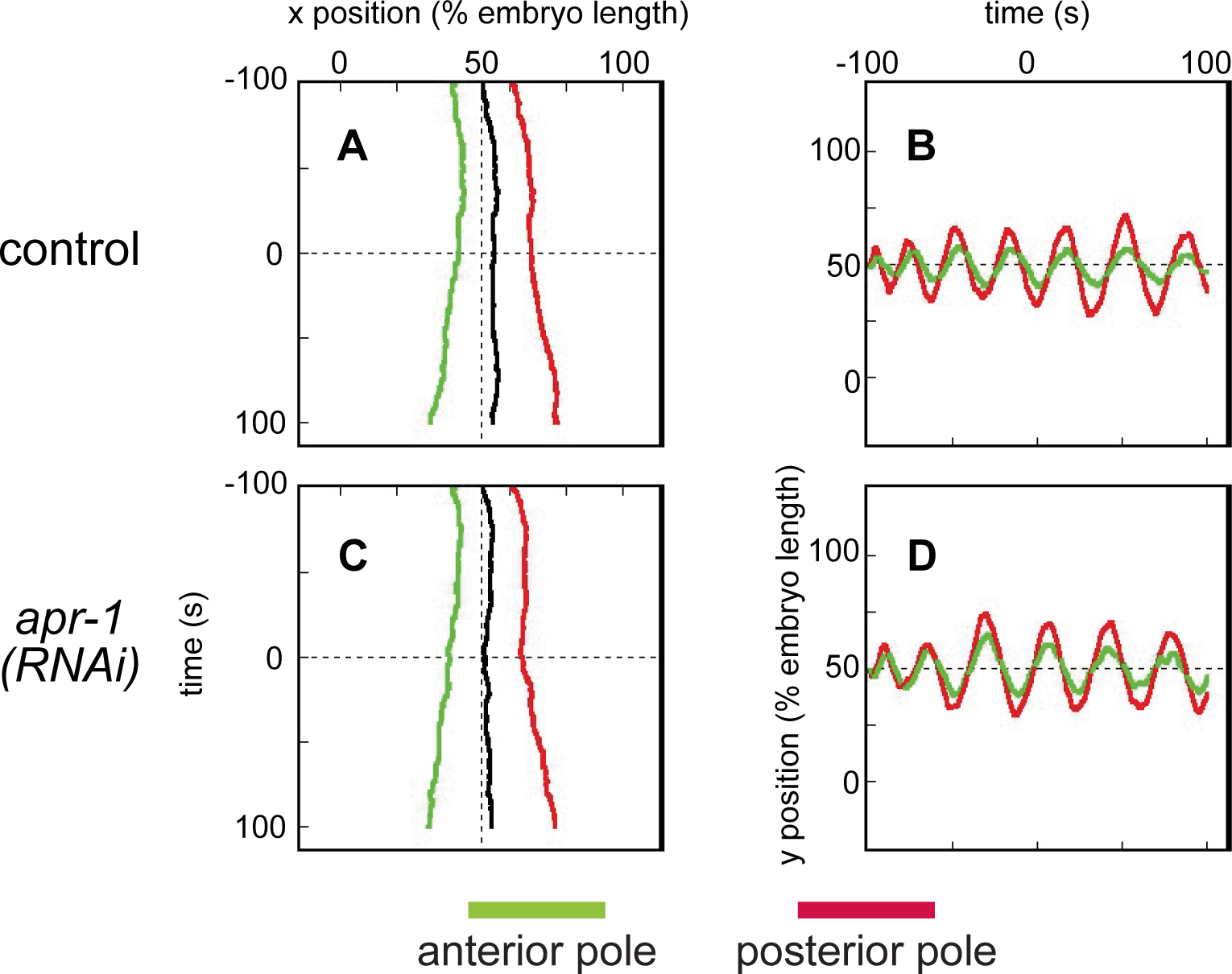
Numerical simulation of spindle movements. (A-D) Representative trajectories of the spindle poles in the simulation. The trajectories of the anterior (green) and posterior (red) poles are shown. Their midpoint (black) is also shown in (A and D). (A, B) Control condition. (C, D) *apr-1(RNAi)* condition. (A and D) Trajectories along A-P axis (x axis). (C and E) and those along an axis perpendicular to the x axis (y axis) are shown.

